# Alternative respiratory electron transport pathways drive differential aminoglycoside susceptibility

**DOI:** 10.64898/2026.05.01.721793

**Authors:** Stuti Srivastav, L. Karvannan, Amitesh Anand

## Abstract

The bacterial electron transport system (ETS) is a highly branched and modular network that interfaces directly with the proton motive force (PMF) to drive ATP synthesis, solute transport, and cellular homeostasis (1–5). Distinct ETS branches differ in their capacities for proton translocation, redox balancing, and membrane polarization. The rewiring of cellular energetics has emerged as a recurrent strategy through which bacteria attenuate drug efficacy by shifting metabolic states, reducing membrane potential, and suppressing reactive oxygen species-linked killing mechanisms (6, 7). Here, we probe the interplay between respiratory routes and antibiotic susceptibilities.

## Result and Discussion

Certain antibiotics enter cells using the PMF driven import system (Figure 1a) (8–10). We tested the hypothesis that the inherent diversity in PMF generation across alternative electron transport branches creates predictable antibiotic vulnerability profiles. *E. coli* has two types of NADH dehydrogenases-(i) proton pumping (two protons per electron) and (ii) non-proton pumping. Also, there are two classes of cytochrome oxidases based on their contributions to PMF generation-(i) cytochrome bo_3_ oxidase (two protons per electron) and (ii) cytochrome bd oxidase (one proton per electron). We engineered four *E. coli* strains with unbranched aerobic ETS by knocking out different combinations of NADH dehydrogenases and cytochrome oxidases (Figure 1b) (11, 12). These strains, hereafter referred to as ETS variants: ETS-1H, ETS-2H, ETS-3H and ETS-4H, should differ in their PMF generation abilities for every electron flowing from NADH to oxygen.

**FIG 1.**
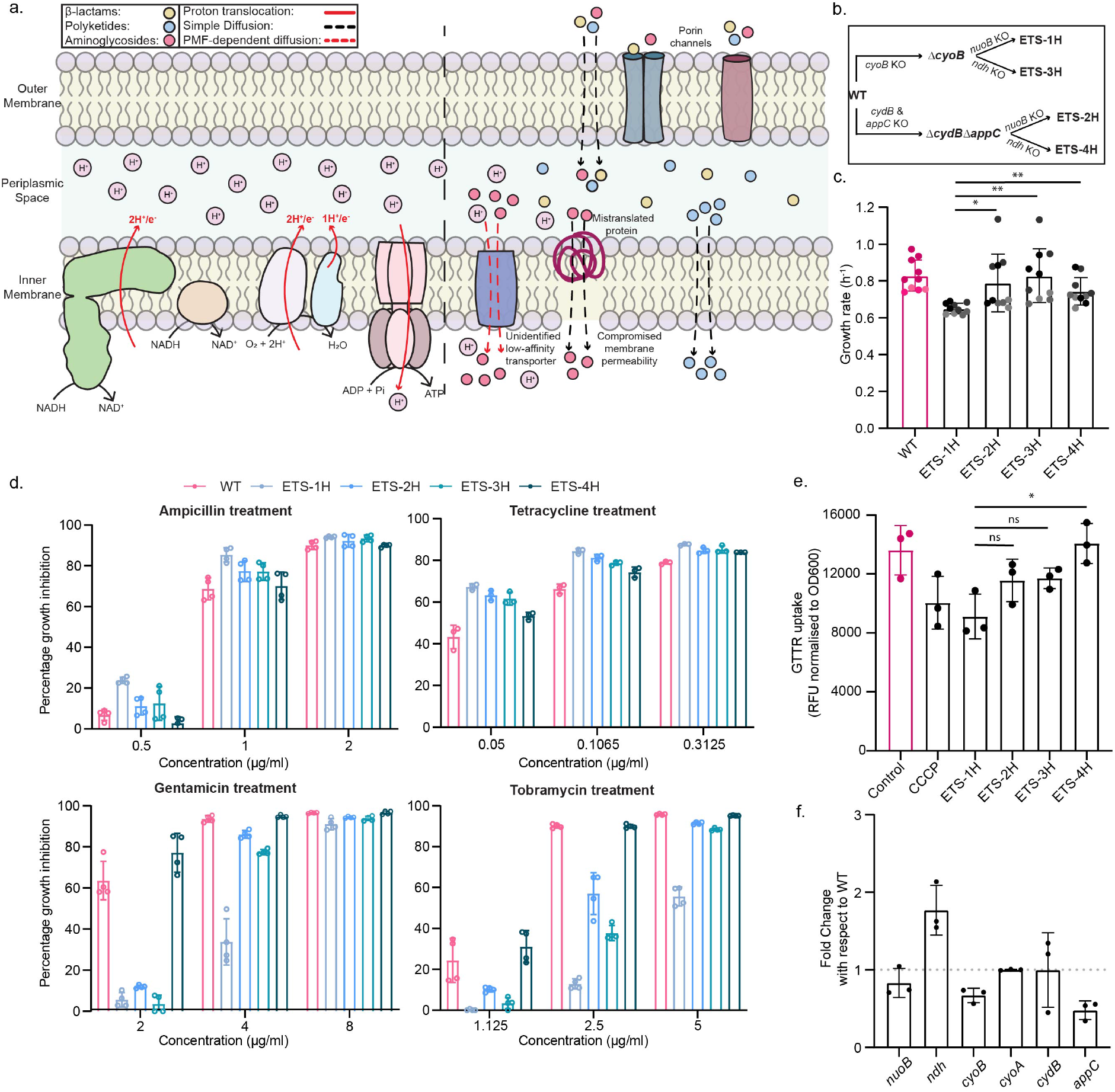
Respiratory chain mutants display differential antibiotic resistance (a) Illustration represents proton translocation during electron transport and modes of entry for different antibiotic classes on the cell membrane of *E. coli*. (b) Schematic for generating ETS variants translocating 1, 2, 3, or 4 proton(s) per electron. (c) Growth rate of the ETS mutants under aerobic conditions. Bar plots represent mean from four biological replicates, each with at least 2 technical replicates, color-coded. Statistics performed using unpaired, two-tailed t-test. (d) Percentage growth inhibition for WT and ETS variants on different antibiotics assessed by the drop in OD600 at 6.5 hours post treatment with a range of antibiotic concentrations, as indicated. (e) Uptake of gentamicin by ETS mutants measured using Gentamicin Texas Red (GTTR) fluorescence. CCCP-treated *E. coli* was used as a positive control. Statistics performed using unpaired, two-tailed t-test. (f) Relative expression of the ETS genes (*nuoB, ndh, cyoB, cyoA, cydB*, and *appC*) in WT strain treated with 2 *µ*g/mL gentamicin with respect to untreated WT (represented by dotted line). Each bar plot represents an average of three biological replicates. Error bars represent standard deviation.

ETS-1H showed a significant growth reduction on M9 minimal medium with glucose as the carbon source (Figure 1c). We performed the antibiotic susceptibility assays for the ETS variants using three different classes of antibiotics: (a) *β*-lactams-Ampicillin, (b) polyketides-Tetracycline, and (c) aminoglycosides-Gentamicin (Figure 1d). We did not observe a significant difference in susceptibility for Ampicillin and Tetracycline among the ETS variants. Remarkably, ETS-1H showed a higher tolerance to Gentamicin as compared to other strains. To probe the class-specific difference in susceptibility, we further examined the tolerance to another aminoglycoside-Tobramycin. We observed a similar higher tolerance in ETS-1H for Tobramycin.

The import of aminoglycosides are reported to be influenced by PMF (9, 13). We probed this relation for the ETS variants using the uptake profile of a fluorescently labeled version of Gentamicin: Texas Red–labeled Gentamicin (GTTR). ETS-1H showed the least fluorescence suggesting lowest Gentamicin uptake (Figure 1e). Notably, the wild-type strain exhibited an uptake profile comparable to ETS-4H, consistent with their similarity in aminoglycoside susceptibility. This observation is puzzling given WT has the option to increase tolerance by shifting to a lower PMF ETS route. There are reports of downregulation of the proton translocating ETS components during the treatment with aminoglycosides-Kanamycin and Gentamicin (14, 15). Thus, there appears to be a metabolic inflexibility in switching ETS route in the given growth condition. While there was an attempt to utilize non-proton-pumping NADH dehydrogenase, we did not observe a clear shift to lower PMF contributory cytochrome oxidases (Figure 1f). Together, our findings establish that architectural choices within the electron transport system are not merely metabolically permissive but actively shape antibiotic vulnerability.

## ACKNOWLEDGMENTS

We thank Mr. Amartya Chowdhury and Ms. Upasana Nandi for their help with validation of some of the data.

## DATA AVAILABILITY STATEMENT

The raw data will be made available upon request

## Materials and method

### Bacterial strains and growth conditions

Experiments were performed using *Escherichia coli* strain C (ATCC 25312) as the wild-type strain. Genes deletions were performed through homologous recombination using templates from the Keio collection (16). P1 phage was used for transduction and recombination (17). The knockouts were confirmed using colony PCR screening. M9 minimal media with 4 g/L of glucose was used for all experiments. Media components were purchased from Sigma-Aldrich. For all the growth assays, absorbance was recorded at optical density of 600 nm using the Tecan Spark multimode microplate reader at 37°C and atmospheric oxygen partial pressure using 96-well plates with 200 *µ*L culture per well. Three different classes of antibiotics were used in this study: *β*-Lactams (Ampicillin), Polyketides (Tetracycline), and Aminoglycosides (Gentamicin and Tobramycin). Tetracycline was dissolved in 70% ethanol and further dilutions were made in autoclaved Milli-Q water (AMQ), while other antibiotics were dissolved directly in AMQ. Growth rate calculations were performed using the gcplyr R package (18).

### Gentamicin-Texas Red (GTTR) uptake assay

*E. coli* WT and mutants were inoculated from glycerol stocks into 2 mL minimal media and allowed to incubate overnight at 37°C and 300 rpm. The optical density at 600 nm of these cultures was noted for normalisation. In 100 *µ*L of overnight grown cultures, carbonyl cyanide m-chlorophenylhydrazone (CCCP; final concentration = 15 *µ*M) and Gentamicin Texas Red (GTTR; final concentration = 2 *µ*g/mL) were added to respective tubes. These were incubated for 30 minutes at 37°C and 300 rpm. The cells were harvested by centrifugation at 15,000 x g for 3 minutes, washed and re-suspended in 100 *µ*L sterile PBS. These were used to detect fluorescence by adding to a white-well microtitre plate. The fluorescence intensities were measured using the Tecan Spark Multimode Plate Reader at excitation wavelength of 595 nm and emission wavelength of 710 nm.

### RNA isolation and qPCR

Bacteria were grown in an M9 minimal medium supplemented with 4 g/L glucose. 2 *µ*g/mL of Gentamicin was added to respective cultures in the mid-exponential phase (OD600 = 0.6 to 0.9). These were incubated at 37°C and 300 rpm for 20 minutes. The culture was harvested and immediately processed for RNA isolation. The cells were dissolved in a TRIzol and incubated at 65°C for 15 minutes. RNA was obtained in an aqueous layer upon the addition of chloroform, which was then precipitated using chilled isopropanol. This RNA pellet was washed with alcohol and finally resuspended in DEPC-treated water. RNA was quantified using a nano spectrophotometer before being frozen at -80°C for later use. cDNA synthesis was performed using Thermo Scientific RevertAid First Strand cDNA synthesis kit using random hexamer primer, according to the manufacturer’s protocol. Quantitative real-time PCR was carried out with BioRad SsoAdvanced Universal SYBR Green Supermix in CFX Opus 96 real-time PCR system (BioRad).

## REFERENCES

1. Biquet-Bisquert A, Carrio B, Meyer N, Fernandes TFD, Abkarian M, Seduk F, Magalon A, Nord AL, Pedaci F. 2024. Spatiotemporal dynamics of the proton motive force on single bacterial cells. Sci Adv 10:eadl5849.

2. Maloney PC, Kashket ER, Wilson TH. 1974. A protonmotive force drives ATP synthesis in bacteria. Proc Natl Acad Sci U S A 71:3896–3900.

3. Strahl H, Hamoen LW. 2010. Membrane potential is important for bacterial cell division. Proc Natl Acad Sci U S A 107:12281–12286.

4. Manson MD, Tedesco PM, Berg HC. 1980. Energetics of flagellar rotation in bacteria. J Mol Biol 138:541–561.

5. Bruni GN, Weekley RA, Dodd BJT, Kralj JM. 2017. Voltage-gated calcium flux mediates mechanosensation. Proc Natl Acad Sci U S A 114:9445–9450.

6. Lobritz MA, Belenky P, Porter CBM, Gutierrez A, Yang JH, Schwarz EG, Dwyer DJ, Khalil AS, Collins JJ. 2015. Antibiotic efficacy is linked to bacterial cellular respiration. Proc Natl Acad Sci U S A 112:8173–8180.

7. Li B, Srivastava S, Shaikh M, Mereddy G, Garcia MR, Chiles EN, Shah A, Ofori-Anyinam B, Chu T-Y, Cheney NJ, McCloskey D, Su X, Yang JH. 2025. Bioenergetic stress potentiates antimicrobial resistance and persistence. Nat Commun 16:5111.

8. Kim S, Eig E, Yue J, Yang A, Comerci CJ, Laune M, Yang C, Kamath A, Shi J, Li P, Cheng Z, Sun C, Guo T, Tian V, Süel GM, Tian B. 2024. Bioelectronic Drug-free Control of Opportunistic Pathogens through Selective Excitability. Device 2.

9. Radlinski LC, Rowe SE, Brzozowski R, Wilkinson AD, Huang R, Eswara P, Conlon BP. 2019. Chemical Induction of Aminoglycoside Uptake Overcomes Antibiotic Tolerance and Resistance in Staphylococcus aureus. Cell Chem Biol 26:1355–1364.e4.

10. Allison KR, Brynildsen MP, Collins JJ. 2011. Metabolite-enabled eradication of bacterial persisters by aminoglycosides. Nature 473:216–220.

11. Anand A, Patel A, Chen K, Olson CA, Phaneuf PV, Lamoureux C, Hefner Y, Szubin R, Feist AM, Palsson BO. 2022. Laboratory evolution of synthetic electron transport system variants reveals a larger metabolic respiratory system and its plasticity. Nature Communications 13:1–9.

12. Patel A, Banwani N, Mink R, Prabhakaran DM, Khairnar SV, Feist AM, Palsson BO, Anand A. 2026. Aerobicity stimulon in revealed using multi-scale computational systems biology of respiratory variants. iScience 29:114715.

13. Webster CM, Shepherd M. 2022. A mini-review: environmental and metabolic factors affecting aminoglycoside efficacy. World J Microbiol Biotechnol 39:7.

14. Choe D, Lee E, Song Y, Kim SC, Jeong KJ, Palsson B, Cho B-K, Cho S. 2025. CRISPRi screening reveals E. coli’s anaerobic-like respiratory adaptations to gentamicin: membrane depolarization by CpxR. mSystems 10.1128/msystems.00353-25.

15. Bie L, Zhang M, Wang J, Fang M, Li L, Xu H, Wang M. 2023. Comparative Analysis of Transcriptomic Response of Escherichia coli K-12 MG1655 to Nine Representative Classes of Antibiotics. Microbiology Spectrum 10.1128/spectrum.00317-23.

16. Baba T, Ara T, Hasegawa M, Takai Y, Okumura Y, Baba M, Datsenko KA, Tomita M, Wanner BL, Mori H. 2006. Construction of Escherichia coli K-12 in-frame, single-gene knockout mutants: the Keio collection. Molecular Systems Biology 2:MSB4100050.

17. Thomason LC, Costantino N, Court DL. 2007. E. coli Genome Manipulation by P1 Transduction. Current Protocols in Molecular Biology 79:1.17.1–1.17.8.

18. Blazanin M. 2024. gcplyr: an R package for microbial growth curve data analysis. BMC Bioinformatics 25:232.

